# Effect of network structure and adaptive foraging on pollination services of species-rich plant-pollinator communities

**DOI:** 10.1101/2024.12.02.626462

**Authors:** Fernanda S. Valdovinos

## Abstract

Network science has had a great impact on ecology by providing tools to characterize the structure of species interactions in communities and evaluate the effect of such network structure on community dynamics. This has been particularly the case for the study of plant-pollinator communities, which has experienced a tremendous growth with the adoption of network analyses. Here, I build on such body of research to evaluate how network structure and adaptive foraging of pollinators affect ecosystem services of plant-pollinator communities. Specifically, I quantify — using model simulations — pollen deposition in networks that exhibit structures like the ones of empirical networks (hereafter empirically connected networks) and those with higher connectance and lower nestedness than empirical networks, for scenarios where pollinators are fixed foragers and scenarios where they are adaptive foragers. I found that empirically connected networks with adaptive foraging exhibit the highest pollen deposition rate. Increased network connectance reduces pollen deposition as increased number of interactions lead to greater conspecific pollen dilution in the absence of other mechanisms such as pollinator floral constancy. High nestedness in moderately connected networks increases the proportion of pollinators visiting only one or two plant species, which are associated with the highest quality visits. Adaptive foraging allows pollinators to quantitatively specialize on specialist plant species which increases conspecific pollen deposition. This research advances pollination biology by elucidating how population dynamics, consumer-resource interactions, adaptive foraging, and network structure affects pollen deposition in a network context.

## Introduction

Network science has substantially contributed to ecology by providing tools to characterize the structure of species interactions within communities and assess how these structures affect community dynamics. This influence is particularly evident in the study of plant-pollinator communities, which have experienced substantial growth due to network analyses. These analyses have enabled much progress in the description of the structure of species interaction within plant-pollinator communities (Bascompte and Jordano 2007, Dormann et al. 2009) and in analyzing the effects of this network structure on the stability of plant-pollinator communities (reviewed in Valdovinos 2019). Such progress, however, has yet to focus on key dynamical and functional properties of plant-pollinator systems beyond their stability, such as quantifying and understanding the mechanisms that determine pollination services.

Pollination services provided by plant-pollinator communities sustain terrestrial biodiversity (Thompson 1994, Ollerton et al. 2011, Ollerton 2017) and food security (Garibaldi et al. 2013, Potts et al. 2016). Despite this key role, it is still unknown how the structure of plant-pollinator networks affects the pollination services these communities provide. Part of the problem are the limitations of Lotka-Volterra type models, which have been widely used to study population dynamics in mutualistic networks (e.g., Bascompte et al. 2006, Bastolla et al. 2009; reviewed Valdovinos 2019) due to their simplicity and mathematical convenience. These models depict mutualistic relationships as net positive effects between species, using a positive term in each mutualist’s growth equation that depends on the partner’s population size. However, by assuming net positive effects phenomenologically, these models cannot quantify the pollination services provided by pollinators to plants nor the effect of network structure on those pollination services. Those mechanisms are confounded together in the net positive effect of the pollinators on the plants.

A more mechanistic alternative to Lotka-Volterra type models is the consumer-resource model by Valdovinos et al. (2013, 2016, 2018). This model breaks down net effects assumed as always positive by Lotka-Volterra models into its biological mechanisms. One of its key innovations is separating plant vegetative dynamics from plant rewards dynamics, allowing: (1) tracking plant reward depletion, (2) assessing exploitative competition among animals visiting the same plants, and (3) incorporating adaptive foraging (behavioral responses to resource availability, Stephens and Krebs 1987, Valdovinos et al. 2010). Another key advancement most related to the study of pollination services is separating the different parts of a plant’s life cycle in processes that affect seed production and those that affect seed recruitment. Interaction with pollinators determine seed production via the quantity and quality of visits a plant receives, which are at the core of the pollination services provided by pollinators. Seed recruitment is determined by plant competition for resources other than pollinators including shared soil resources and light, which is modeled more phenomenologically.

I use the Valdovinos et al’s dynamic model to test three hypotheses (Fig. 1) on how network structure and adaptive foraging affects pollination services, which I quantify as pollen deposition rate. First, I hypothesize that increasing network connectance (i.e., the fraction of potential interactions that are realized) decreases pollen deposition rate. This because higher connectance results in pollinators visiting more plant species, which — without other mechanisms in place (e.g., pollinator floral constancy) — will increase conspecific pollen dilution (Ne’eman et al. 2010, Brosi 2016). Second, I hypothesize that nestedness (i.e., the tendency of generalist species to interact with both specialist and generalist species, and specialist species to interact with only generalist species) will decrease pollen deposition rate in networks without adaptive foraging by increasing niche overlap among plant species for pollination services of shared pollinator species (Fig. 1A). Third, I hypothesize that adaptive foraging will increase pollen deposition rate by allowing niche partitioning among plant and pollinator species (Valdovinos et al 2013, 2016), which will reduce conspecific pollen dilution (Fig. 1B).

**Figure 1.**
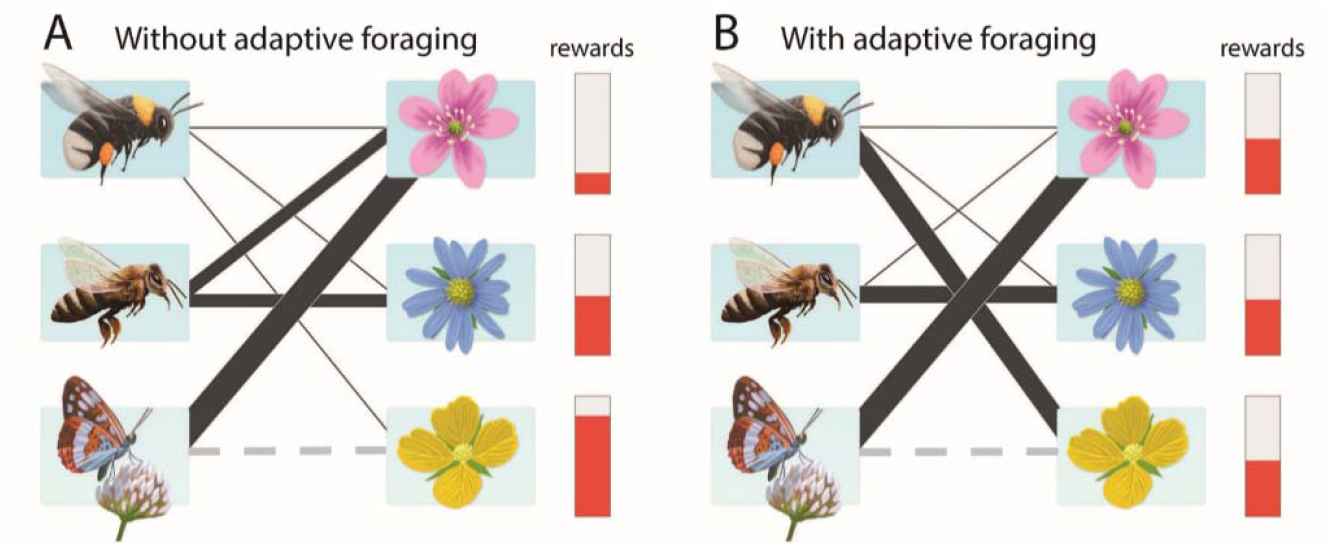
Rational behind my three hypotheses. **(A)** Illustration of the behavior of the model without adaptive foraging (ignore gray dashed line) where pollinators assign the same fixed foraging effort to each of the plant species they visit (illustrated by the thickness of the links connecting each pollinator species to each plant species in its diet), which results in generalist plants receiving more and higher quality of visits and having reduced floral rewards (red bars) than specialist plants. **(B)** Illustration of the behavior of the model with adaptive foraging (AF) where pollinators reassign more foraging effort to the plant species in their diet that have higher floral rewards available (specialist plant species), which results in specialist plants receiving more and higher quality of visits than without adaptive foraging. Note that **(A)** also represents the initial state for networks with adaptive foraging before the generalist pollinators reassign their foraging efforts to specialist plants, and that both **(A)** and **(B)** represent nested networks (ignore dashed gray line). My first hypothesis is that increasing connectance, illustrated by adding a new interaction (gray dashed line), decreases overall pollen deposition rate in the network by decreasing pollinator visit quality. For example, the added interaction in **(A)** results in the quality of visits of the specialist pollinator (bottom-left) assigned to the generalist plant (top-right) from σ_*ij*_ = 1 to σ_*ij*_ = 0.5 (see Methods). Nestedness increases the fraction of interactions shared by generalist (top) and specialist (bottom) species. For example, all three plant species in both **(A)** and **(B)** share the pollination services performed by the most generalist pollinator species, the bumblebee. Thus, I hypothesize that nestedness decreases pollen deposition rate by increasing dilution of conspecific pollen carried by generalist pollinators. With adaptive foraging **(B)**, generalist pollinators assign higher foraging effort on specialist plants (follow thick line) which increases the quantity and quality of visits performed by generalist pollinators on specialist plants, while specialist pollinators keep assigning all their visits to generalist plants. Therefore, I hypothesize that adaptive foraging increases overall pollen deposition rate.

## Methods

The Valdovinos et al’s model (Valdovinos et al. 2013, 2016, Valdovinos and Marsland 2020) defines the population dynamics of each plant (Eq. 1) and pollinator species (Eq. 2) of the network, as well as the dynamics of floral rewards (Eq. 3) of each plant species, and the foraging effort (Eq. 4) that each pollinator species (per-capita) assigns to each plant species in its diet. Note that α_*ij*_ = 0 if pollinator *j* does not visit plant *i*, that is, α_*ij*_ encapsulates the plant-pollinator network. Important to the objective of quantifying pollination services is that this model calculates the quantity and quality of visits received and performed by each plant and pollinator species, and the resulting pollen deposition rate. Table 1 lists all the variables and parameters of the model, their definition, values, and units. Table 2 lists the biological mechanisms in the model with their assumptions. The governing equations of this model are as follows:

**Table 1.**
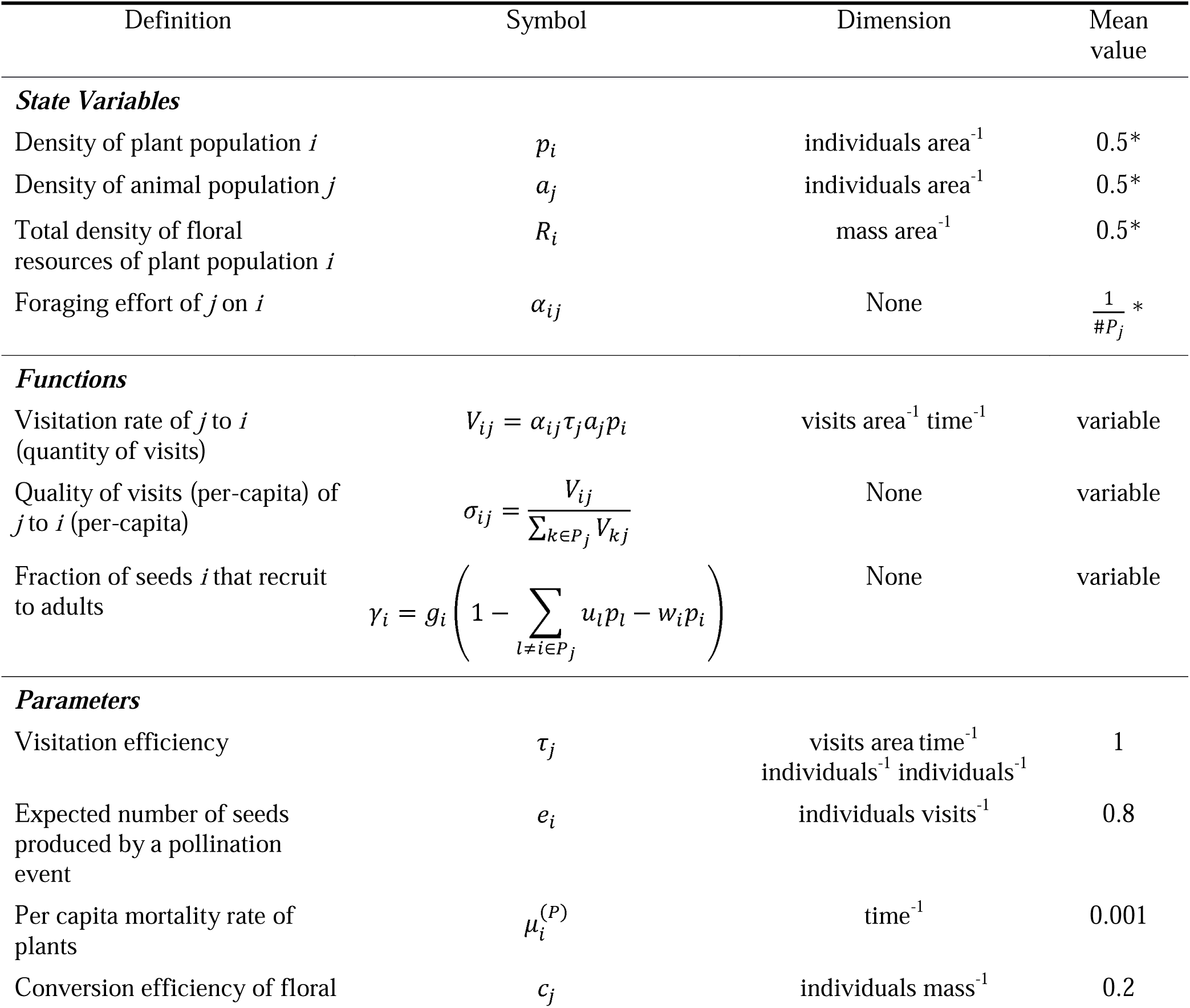
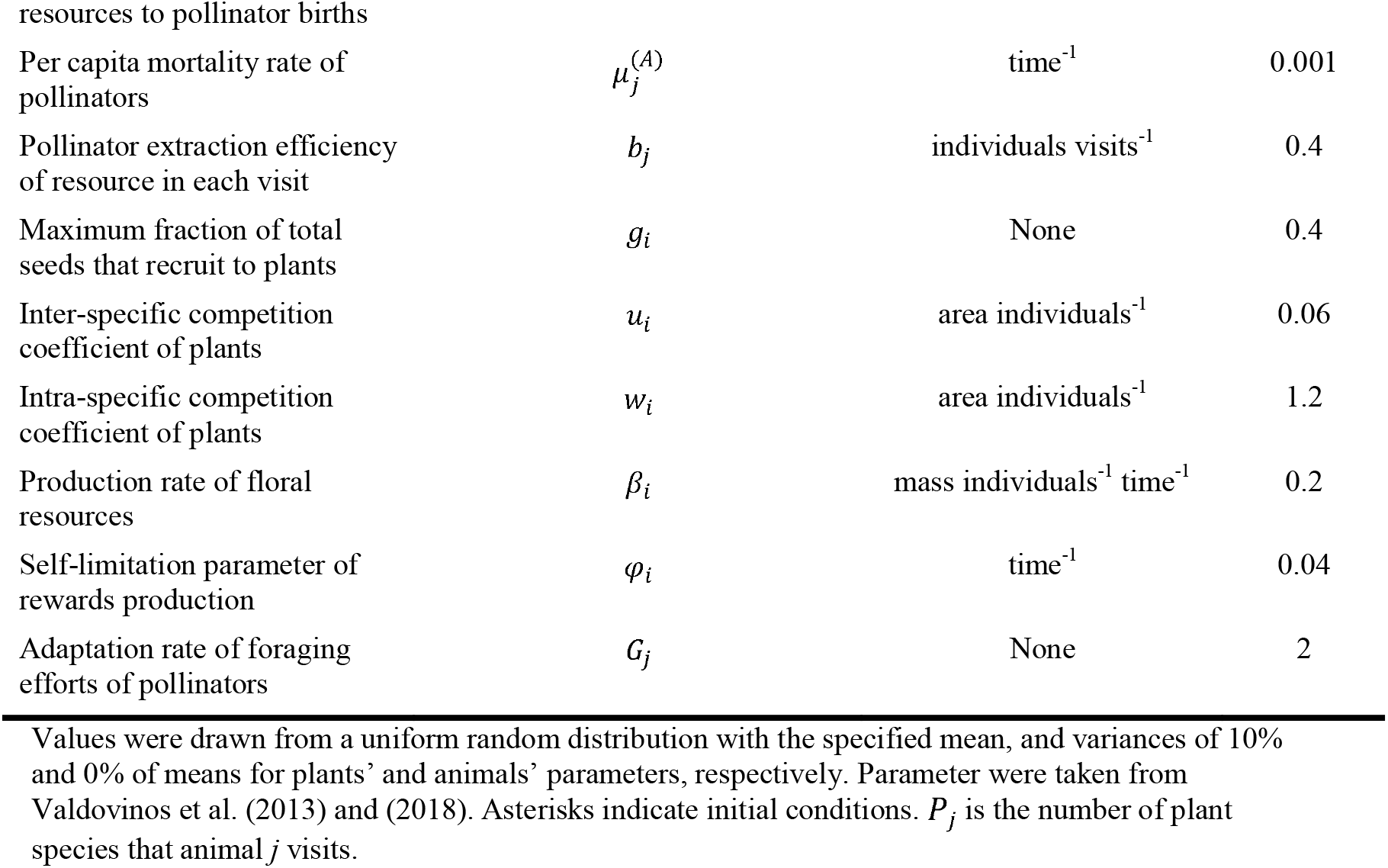
Model state variables, functions, and parameters.

**Table 2.**
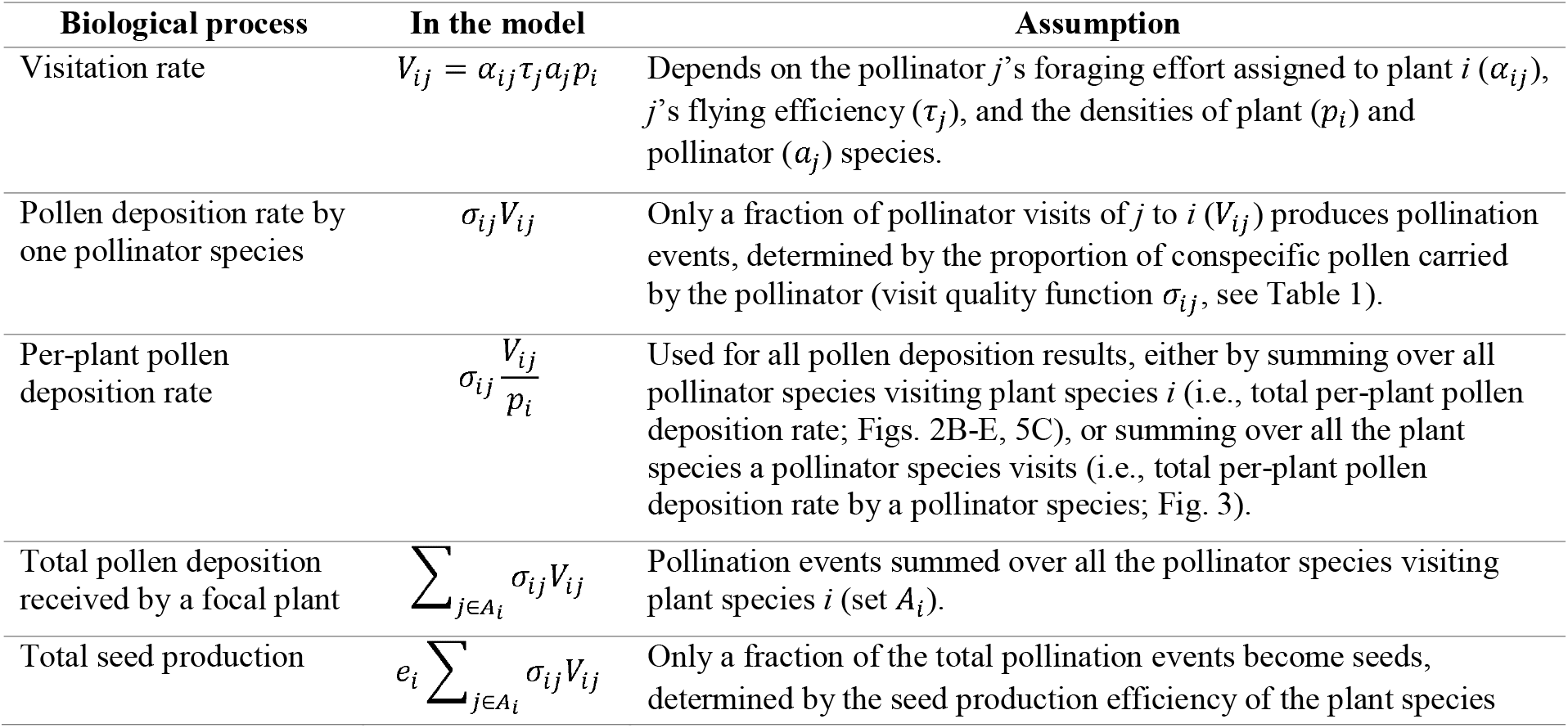
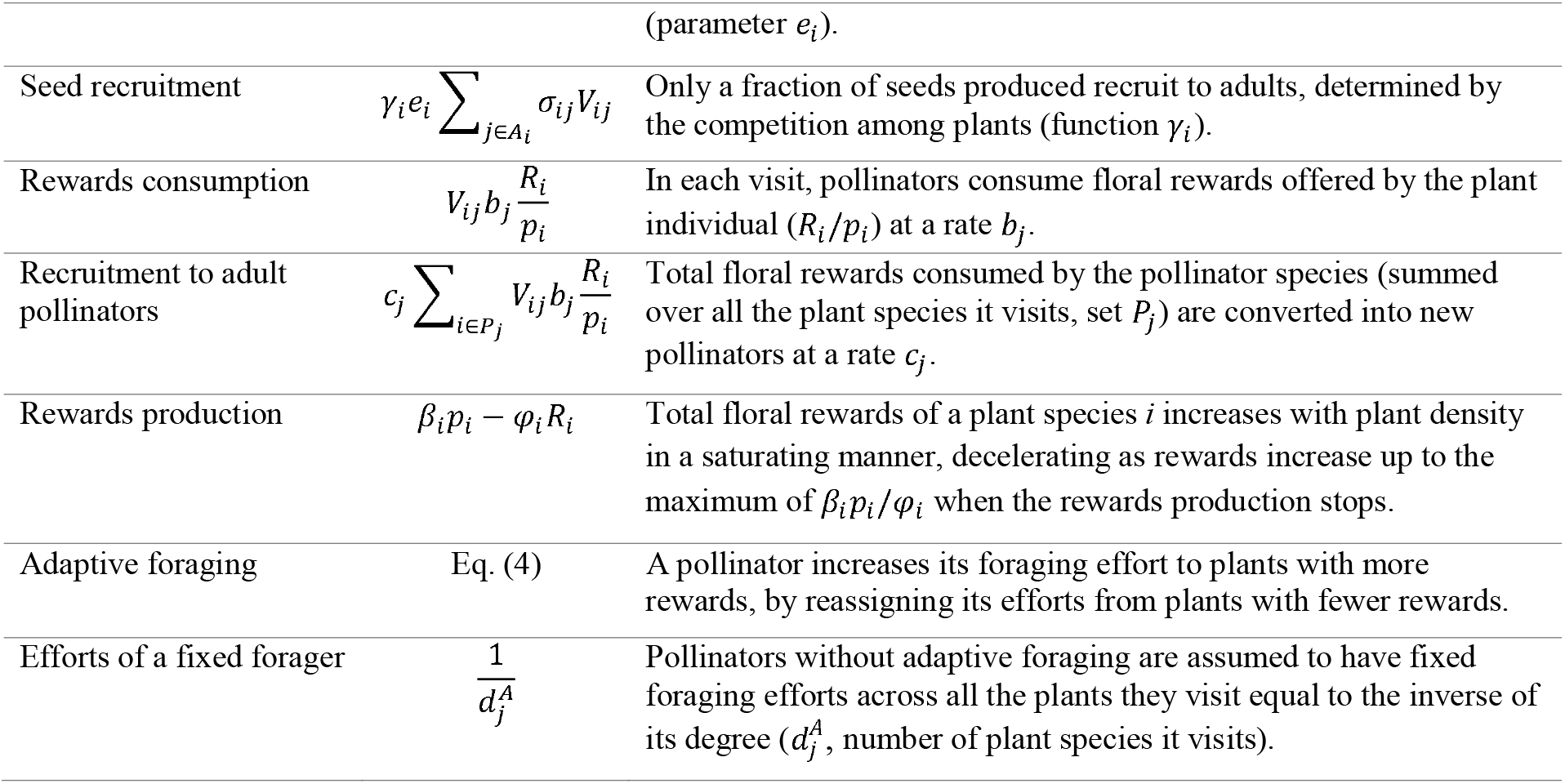
Biological processes, assumptions, and key dynamic quantities in Valdovinos et al.’s (2013) model, and plant coexistence metrics.

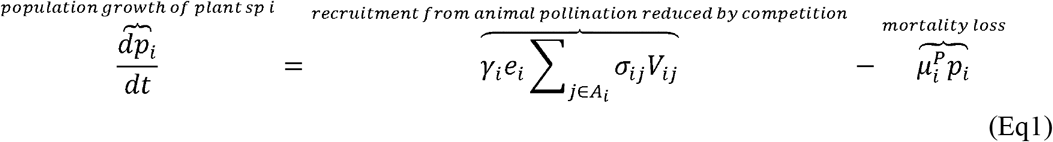

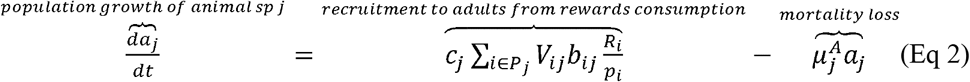

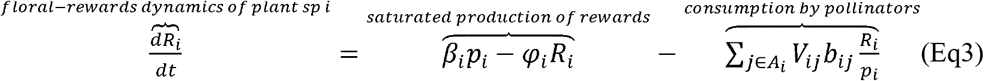

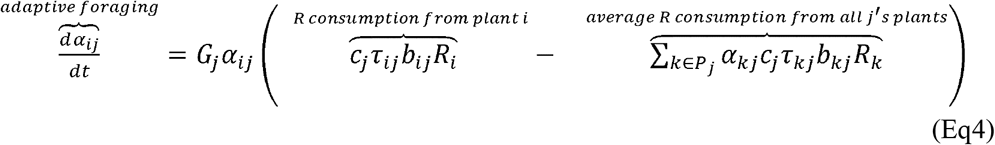

I applied this model to 800 networks generated by a widely used algorithm (Thébault and Fontaine 2010) that allows manipulating the networks’ species richness, connectance, and nestedness. I generated 100 networks to have species richness (*S*) and connectance (*C*) centered at S=90 and C=0.15, S=90 and C=0.30, S=200 and C=0.15, and S=200 and C=0.30, half non-nested and the other half nested (Fig. 2A). That is, 100 networks per each of the 8 network-structure treatments (2 richness levels × 2 connectance levels × 2 nestedness levels). Only the 100 nested networks centered at S=90 and C=0.15, are empirically connected (noted by EC in Fig. 2B), that is, exhibit structures like the ones of empirical networks (Fig. 2A). I run the model for each of the 800 networks without and with adaptive foraging. Pollinators in networks without adaptive foraging assign fixed and equal foraging efforts to each of the plant species in their diet

**Figure 2.**
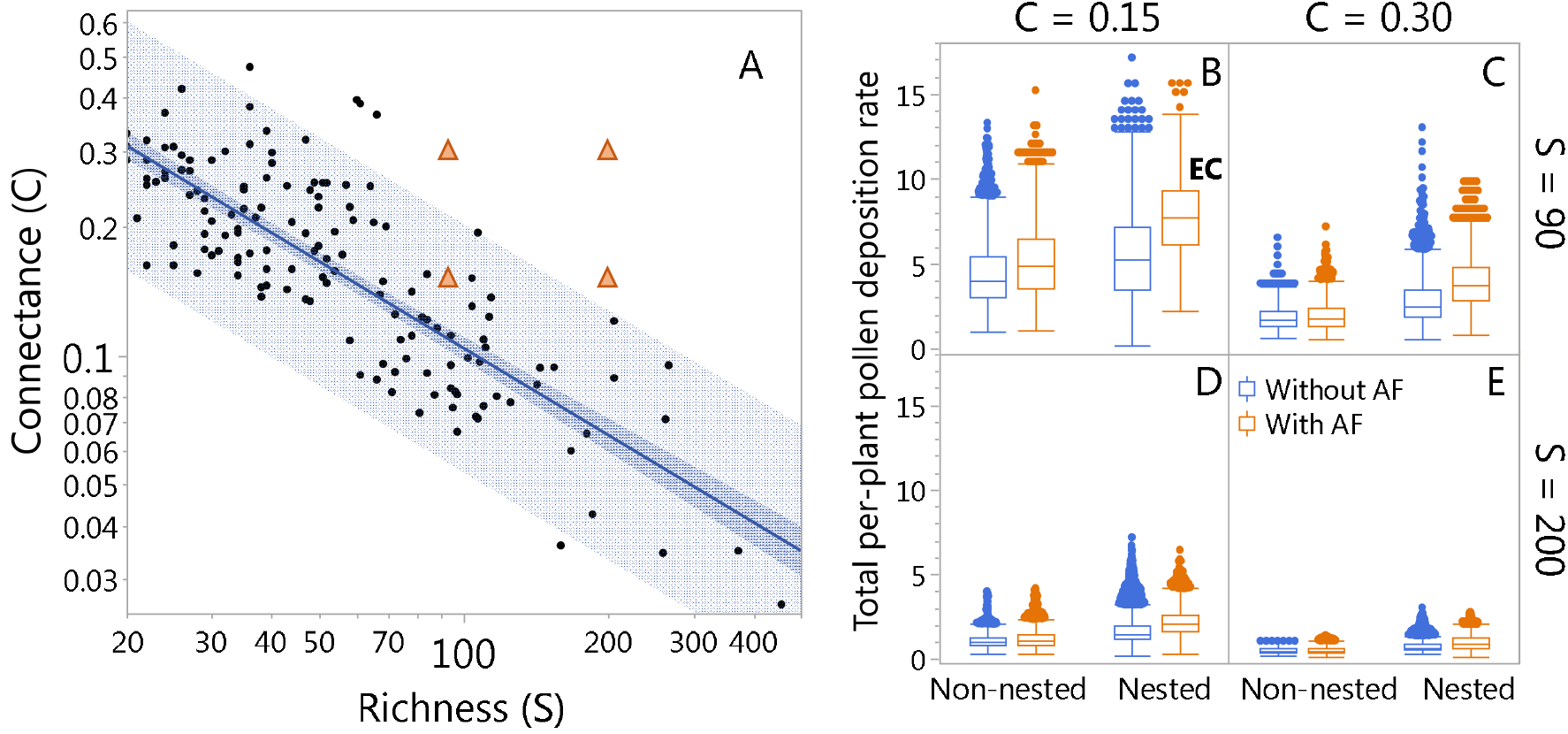
Effects of network structure and adaptive foraging on pollen deposition rate. Panel **A** shows connectance (C) vs species richness (S) of 172 empirical plant-pollinator networks obtained from https://www.web-of-life.es/ that contain between 15 and 500 species. Dots represent the empirical networks. The blue solid line and darker shaded area represent the regression between C and S and its confidence interval, respectively. The lighter shaded area is the 95% prediction interval. The orange triangles represent the C and S at where the algorithm-generated networks were centered, that is S=90 and C=0.15, S=90 and C=0.30, S=200 and C=0.15, and S=200 and C=0.30. The other four panels show the pollen deposition rate summed over all visits received by plant species on a per-capita basis (see Table 2), for networks centered at S=90 (**B, C**) and S=200 (**C, D**) with connectance centered at C=0.15 (**B, D**) and C=0.30 (**C, E**), each at two levels of nestedness (non-nested and nested), totaling 800 networks, without (blue) and with (orange) adaptive foraging (AF). Boxes indicate first and third quantile with middle line representing the median, and error bars max and min values without outliers. Dots represent outliers. EC stands for empirically connected networks (in this case, nested networks with S=90 and C=0.15).

## Results

Among all 8 network-structure treatments of 100 networks each, the empirically connected (i.e., nested with S=90 and C=0.15; EC in Fig. 2B) exhibit the highest per-capita pollen deposition rate. Increasing connectance beyond what it is observed in empirical networks reduces pollen deposition rate in all richness and nestedness levels consistently with my first hypothesis. The mechanism explaining the negative effect of increased connectance on pollen deposition rate rests in the number of specialist pollinator species in a network. Pollinators visiting fewer plant species are the most efficient in depositing conspecific pollen (Fig. 3) with specialist pollinators (those visiting only one plant species) being the most efficient of all pollinators. The more overconnected the networks are with respect of what it is observed in empirical networks (Fig. 2A), the fewer the specialist species in the network (Figs. 3, 4E-H). For example, the most overconnected treatments (i.e., the 200 non-nested and nested networks centered at S=200 and C=0.3) have zero specialist pollinator species with the only exception of one nested network that has one specialist pollinator species.

**Figure 3.**
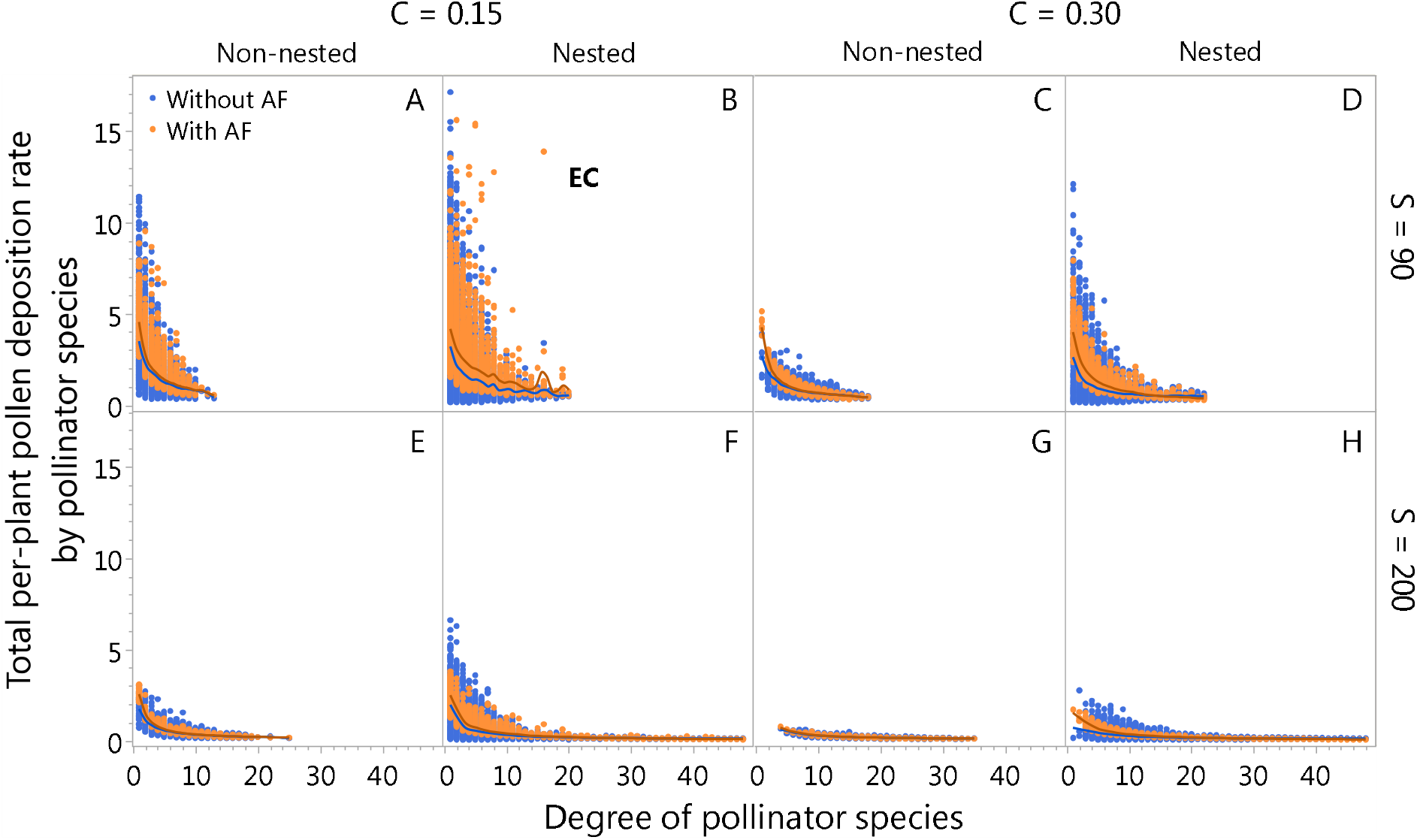
Pollinator species visiting the fewest plant species are the most efficient pollinators in terms of pollen deposition rate. Dots represent the total per-plant pollen deposition rate by each pollinator species in each of the 800 networks, without (blue) and with (orange) adaptive foraging (AF). Panels A-D show results for networks with species richness centered at S=90, while panels E-H show results for networks with richness centered at S=200. Panels A, B, E, F show results for networks with connectance centered at C=0.15, while panels C, D, G, H show results for networks with connectance centered at C=0.3. Panels A, E, C, G show results for non-nested networks, while panels B, F, D, H show results for nested networks. Panel B shows the empirically connected (EC) networks, which exhibit the highest pollen deposition rate by specialist pollinators. Conversely, the more overconnected the networks are with respect of what it is observed in empirical networks (Fig. 2A), the fewer the specialist species with none in non-nested networks with species richness centered at S=90 and connectance at C=0.3 (see Fig. 4D).

**Figure 4.**
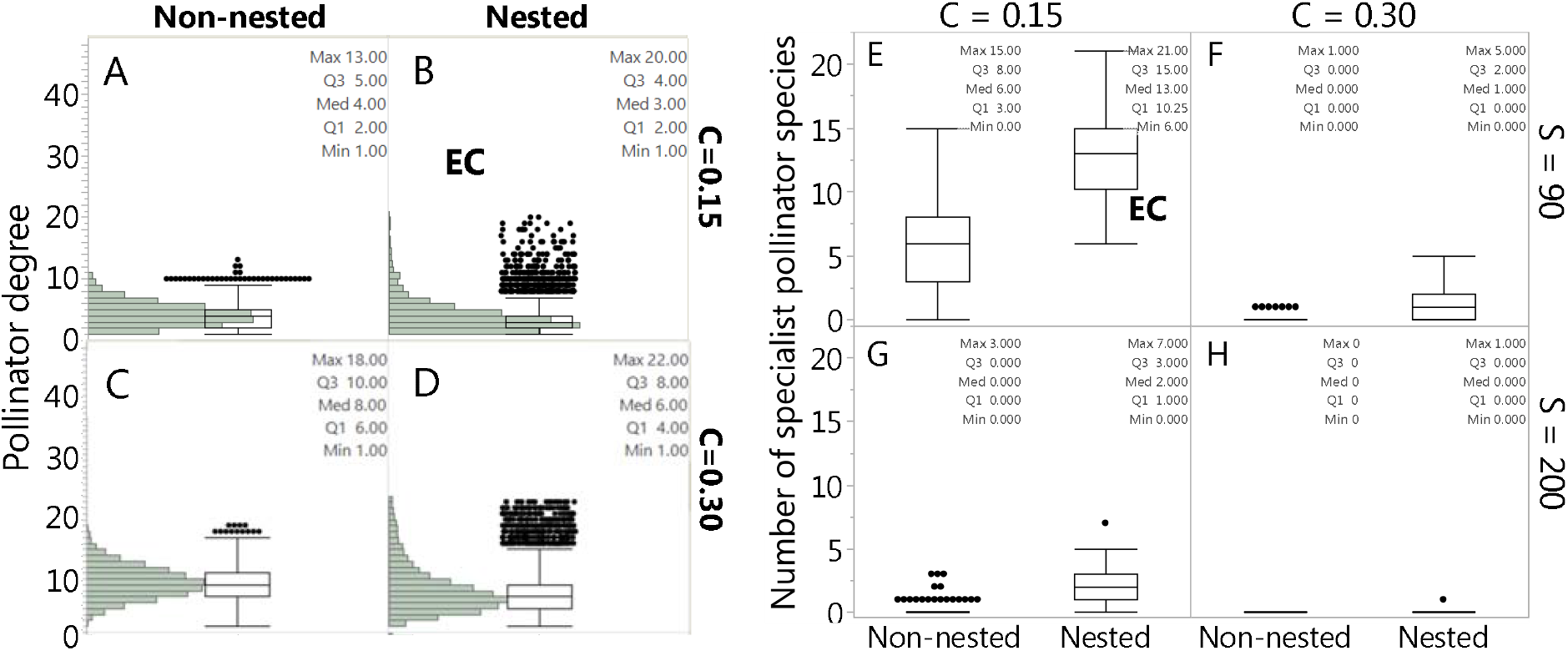
Pollinator degree distribution and number of specialist pollinator species in simulated networks. Panels (A-D) show histograms for the frequency of pollinator degrees (i.e., number of plant species each pollinator species visits) across all networks with species richness centered at S=90, separated by nestedness (non-nested and nested) and connectance (C=0.15 and C=0.3) levels. Panels (E-H) show the number of specialist pollinator species (i.e., those visiting only one plant species) in all 8 network-structure treatments, separated by species richness (S=90 and S=200), connectance (C=0.15 and C=0.3) and nestedness (non-nested and nested) levels. Note that the number of specialist species from panels A and B (i.e., degree 1) are both plotted in panel E, while the number of specialist pollinators from panels C and D are plotted in panel F. The degree distributions of networks with species richness centered at S=200 are not shown due to space limitations, but they look as normal distributions like panel C. Text inside every panel indicate the maximum (Max), median (Med), minimum (Min), and the first (Q1) and third (Q3) quartiles.

Contrary to my third hypothesis, nestedness increases pollen deposition rate. This positive effect of nestedness is explained by the positive correlation between nestedness and the heterogeneity of degree distribution. That is, more nested networks exhibit more skewed degree distributions (Figure 4A-D) with most species having few interactions while few species monopolizing most of the interactions in the network. Figure 4A-D show histograms for pollinator degree distributions showing different levels of skewness for networks with species richness centered at S=90 depending on the network-structure treatment. The degree distribution of the empirically connected networks (nested with C=0.15; Fig. 4B) is the most skewed, with a median of degree 3, a high frequency of degree 1, and a long-tail of very low frequencies of higher degrees reaching a maximum of degree 20. In contrast, non-nested networks with the same connectance (C=0.15) and species richness (S=90) exhibit a minimal tail with hyper-generalists visiting only up to 13 plant species (Fig. 4A). More importantly for increasing pollen deposition rate in the model, nestedness increases the proportion of specialist pollinators that visit only one plant species (compare the number of specialist pollinators in non-nested and nested networks in Fig 4E) which are the pollinators performing the highest pollen deposition rate (Fig. 3).

As mentioned above, the empirically connected (EC, Fig. 4E) have the largest number of specialist pollinators among the simulated networks, with a median of 13 (minimum and maximum of 6 and 21, respectively). Empirical networks, however, exhibit even higher number of specialist pollinator species (Fig. 5) than the EC networks. At the species richness range comparable with the simulated networks (i.e., S=88-205 in Fig. 5), the empirical networks have a median of 33 pollinator species (with minimum and maximum of 6 and 103, respectively). Considering that the number of specialist pollinator species increases with species richness (Fig. 5), even empirical networks with fewer species than the EC networks (see S=54-88) have more specialist pollinator species, with a median of 19 (minimum of 0 and maximum of 36).

**Figure 5.**
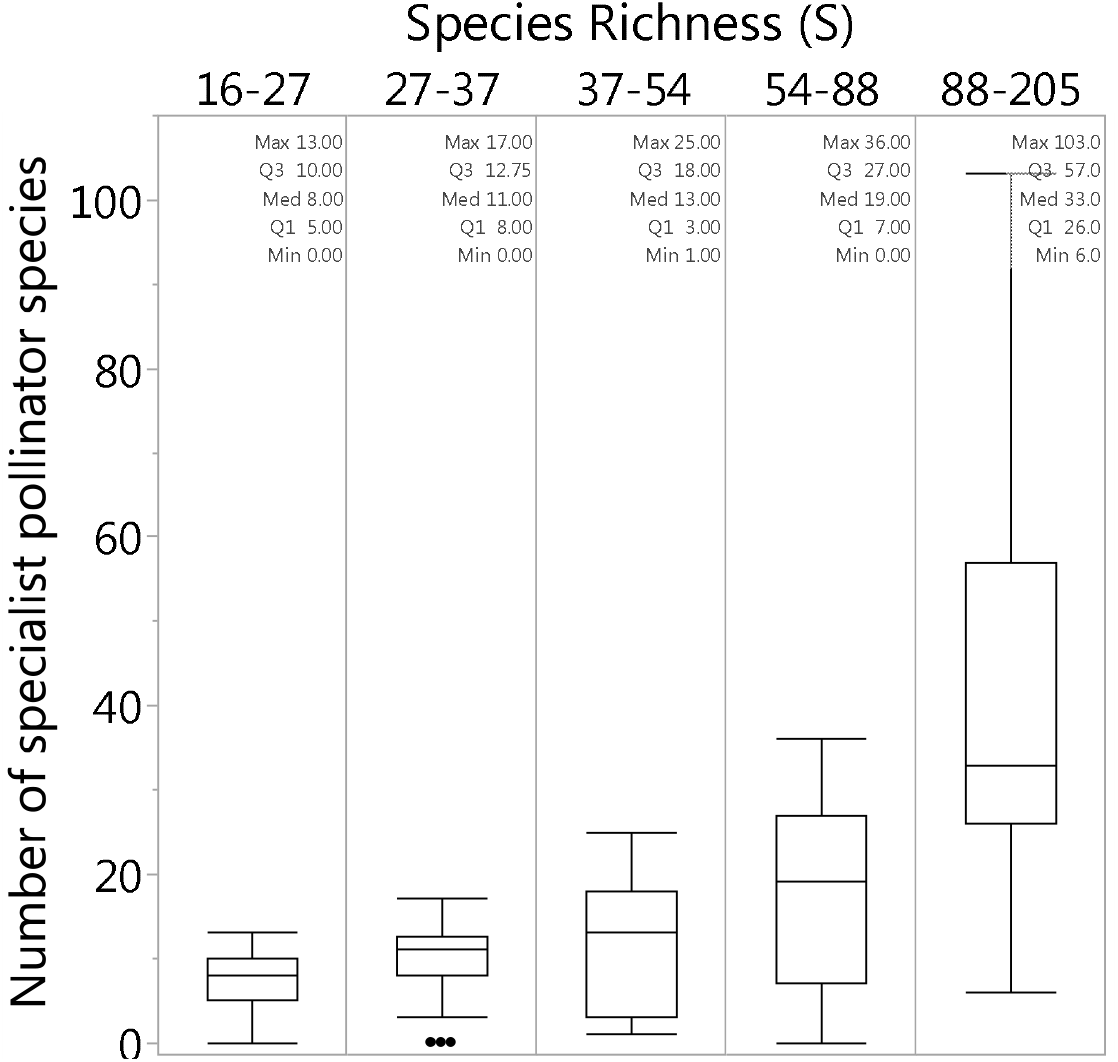
Number of specialist pollinator species in empirical networks. Empirical networks have more specialist pollinator species than the simulated networks, number that increases with species richness. At the species richness range comparable with the empirically connected networks (i.e., S=88-205), the empirical networks have a median of 33 pollinator species (with minimum and maximum of 6 and 103, respectively). Boxes indicate first and third quantile with middle line representing the median, and error bars max and min values without outliers. Dots represent outliers.

Confirming my third hypothesis, adaptive foraging increases pollen deposition rate but only slightly and mostly in EC networks (Fig. 2B). A clearer result of adaptive foraging on pollen deposition, however, is in inverting its relationship with plant degree from positive to negative (Fig. 6A). That is, adding adaptive foraging to the network dynamics results in specialist plants experiencing higher pollen deposition rates than generalist plants, while generalist plants exhibit higher pollen deposition rates than specialists in networks with fixed foragers. Figure 6B shows that in networks without AF, pollinators assign the same foraging efforts 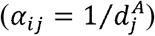 to each of its plant species regardless of plant degree. Figure 6C shows that pollinators with adaptive foraging re-assign their foraging efforts to plant species in their diet that have lower degree. Thus, the positive effect of adaptive foraging on pollen deposition of specialist plants occurs due to pollinators re-assigning most of their foraging effort from the generalist to the specialist plant species in their diet (compare Figs 6B and 6C).

**Figure 6.**
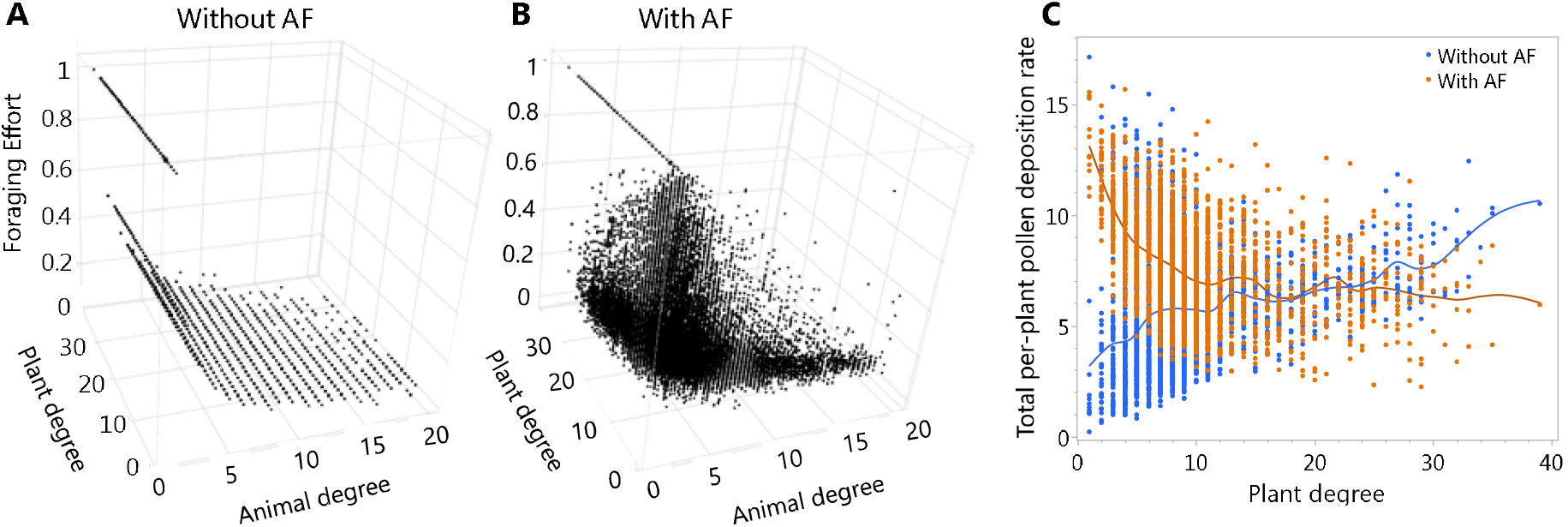
Adaptive pollinators quantitatively specialize on specialist plant species which increases pollen deposition for specialist plants and decreases pollen deposition for generalist plants. Foraging efforts that pollinators assign to plants with respect of the degree of the pollinator and plant species interacting in networks without (A) and with (B) adaptive foraging (AF). **A)** Without AF, all pollinator species with the same degree 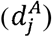 have the same fixed foraging effort of 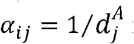 for each plant species in their diet regardless of the degree of the plant species, equal to. **B)** With AF, pollinators quantitatively reassign their foraging efforts from generalist to specialist plants. **C)** Total per-plant pollen deposition rate with respect of plant degree without (blue) and with (orange) adaptive foraging (AF). Note that the positive relationship between pollen deposition rate and plant degree without AF is reversed with AF. That is, the preference of adaptive pollinators for specialist plants results in specialist plant species having a higher per-plant pollen deposition rate than generalist plant species in networks with AF, while the inverse happens in networks without AF. The line of points at pollinator degree 1 in both Figs. 6B and 6C, correspond to specialist pollinator species, whose foraging effort and quality of visit stay at its maximum of 1, which is assigned to the only plant species in their diet.

## Discussion

Among the eight network-structure treatments, empirically connected networks exhibit the highest per-capita pollen deposition rate. Increasing connectance beyond the levels observed in empirical networks reduces pollen deposition rates across all levels of species richness and nestedness, consistent with my first hypothesis. In contrast, nested networks show higher pollen deposition rates than non-nested networks, contradicting my second hypothesis. Finally, adaptive foraging slightly increases pollen deposition rates, supporting my third hypothesis, and reverses the positive relationship between pollen deposition rate and plant degree.

The effect of network structure—specifically connectance and nestedness—on pollen deposition rates is driven by the proportion of specialist pollinators within the network. Networks with a higher proportion of specialist pollinator species exhibit greater pollen deposition rates. The simulated networks that were moderately connected and highly nested (i.e., the empirically connected networks) exhibited the highest pollen deposition rates among all simulated networks due to having the highest proportion of specialist pollinator species. To evaluate whether the number of specialist pollinators in my simulated networks aligns with empirical data, I compared it to empirical networks and found that the latter contained even higher proportions of specialist pollinators. However, the true existence of such a high number of specialist pollinator species in empirical networks has been questioned (Waser et al. 1996, Brosi 2016; but see Johnson and Steiner 2000), as it may result from insufficient sampling effort rather than reflecting true ecological patterns (Blüthgen et al. 2008, Blüthgen 2010).

This questioning has prompted numerous studies investigating the effects of sampling effort on network structure, both through field studies (Nielsen and Bascompte 2007, Hegland et al. 2010, Chacoff et al. 2012) and through models generating network structures (Blüthgen et al. 2008, Blüthgen 2010, Fründ et al. 2016). These studies have consistently shown that incomplete sampling strongly underestimates the number of interactions while simultaneously overestimating the degree of specialization. However, for the present results, pollinator specialization does not need to occur at the species level for its positive effects on pollen deposition rates to hold. Instead, specialization can occur at the individual level through flower constancy (Chittka et al. 1999) or fidelity (Brosi 2016), or as temporary, realized specialization at the population level. Flower constancy (or fidelity more broadly) is the tendency of pollinators to visit flowers of the same species during a foraging bout despite the presence of other available floral resources. This behavior allows individual pollinators to act as specialists during their foraging bouts, even if the species as a whole interacts with multiple plant species. By focusing on a single plant species throughout a foraging bout, individual pollinators increase the efficiency of pollen transfer and deposition (Chittka et al. 1999, Ne’eman et al. 2010, Brosi 2016), producing results similar to those presented here.

Another mechanism that could produce results similar to those presented here is realized specialization. This term refers to pollinator species that are observed visiting only one plant species during a specific sampling period, even though they may interact with other plant species at different times or under varying conditions. These pollinators are not true specialists in an evolutionary sense but instead exhibit flexibility in their interactions, often through turnover or rewiring over time (Brosi 2016, CaraDonna et al. 2017, Glaum et al. 2021). To summarize, any mechanism of specialization—whether temporary, individual-level, population-level, or species-level—suffice for the present results of increased pollen deposition to hold.

Beyond the number of specialist pollinators, higher connectance increases the number of plant species each pollinator interacts with. This broader range of interactions per pollinator species can lead to heterospecific pollen transfer (Arceo-Gómez et al. 2020) and dulition in conspecific pollen on pollinators’ bodies. Nestedness, however, counteracts these effects by creating a skewed degree distribution among plant and pollinator species. In highly nested networks, the degree distribution follows a long-tail pattern (Payrató-Borras et al. 2019), where most species have few interactions, and a small number of species dominate most interactions within the network. Therefore, nested networks will always have many pollinator species with one or few realized interactions which will be effective pollinators in terms of pollen deposition.

Adaptive foraging further enhances pollen deposition rates in moderately connected and highly nested networks by allowing pollinators to allocate greater per-capita foraging efforts to specialist plants. Specialist plants, therefore, benefit from receiving higher-quality visits from generalist pollinators that focus their efforts on specialist plants and primarily carry their conspecific pollen (Valdovinos et al. 2013, 2016, Valdovinos and Marsland 2020). This occurs because specialist plants in the model tend to have more available rewards, as they are less frequently visited compared to generalist plants, which experience higher visitation rates (Valdovinos et al. 2013, 2016, Valdovinos and Marsland 2020).

In conclusion, this study demonstrates that the structure of plant-pollinator networks and the adaptive foraging behavior of pollinators can play critical roles in determining pollen deposition rates in plant-pollinator communities. Empirically connected networks with moderate connectance and high nestedness exhibit the highest pollen deposition rates, highlighting the importance of network structure in facilitating effective pollination. While increased connectance can dilute conspecific pollen through heterospecific interactions, nestedness counteracts this effect by creating a degree distribution that promotes specialization, even if only temporarily or at the individual level. Adaptive foraging further enhances pollen deposition by enabling pollinators to allocate more effort to specialist plants, thereby improving resource partitioning and coexistence between specialist and generalist species.

These findings emphasize the interplay between network structure, pollinator behavior, and ecosystem function, advancing our understanding of how plant-pollinator communities sustain critical ecological processes. Future research should more explicitly evaluate the interplay between individual pollinator specialization and network structure, which can be studied both empirically and by using mathematical models. By integrating network theory, behavioral ecology, and ecosystem dynamics, this work provides a framework for understanding the complex relationships that underpin ecosystem services such as pollen deposition.

## Data Availability Statement

The code used to produce all results of this work is publicly available at GitHub repository https://github.com/Valdovinos-Lab/PollenDeposition.

## Acknowledgements

This work was funded by National Science Foundation (NSF) gran DEB-2129757.

